# Age-related changes in the gut microbiota of a long-lived seabird suggest divergence from mammalian models

**DOI:** 10.1101/2025.10.20.683453

**Authors:** Eveliina Hanski, Kieran A Bates, Robert Hughes, Mark Newell, Alex Penn, Aura Raulo, Liz Scott, Liping Tang, Annette L Fayet

## Abstract

Gut microbiota plays key roles in shaping host development, immunity, and physiology. The assembly of vertebrate gut microbiota during early life follows relatively predictable trajectories across several mammalian species, largely driven by maternal transmission routes. However, birds lack these transmission mechanisms due to their oviparity, potentially leading to different age-related microbiota patterns. Here, we investigate gut microbiota variation in Atlantic puffin (*Fratercula arctica*) chicks and adults across six British populations using 16S rRNA amplicon sequencing. We find that chick microbiotas show strong population-level structuring that disappears in adults, with spatial patterns particularly pronounced in anaerobic bacteria. Puffin microbiotas become increasingly individualised and aerotolerant with age, while taxonomic richness remains stable – patterns that contrast with the convergent, richness-increasing trajectories observed in several mammalian species. Our results suggest that environmental exposure drives early-life microbiota assembly in birds, highlighting the value of avian systems for understanding vertebrate microbiota development beyond mammals.

## Introduction

The vertebrate gut microbiota plays essential roles in shaping host physiology, development, and immune function^1–3^. At the same time, gut microbial communities are highly dynamic, varying across hosts, populations, life-stages, and environments^4–10^. Understanding the ecological and developmental drivers of this variation is key to linking microbiota structure with function in natural systems.

Most insights into gut microbiota dynamics come from studies in humans and laboratory animals^1,11–13^, with wild mammals receiving increased attention in recent years^14–19^. Despite birds accounting for >5,000 more species than mammals^20^, avian gut microbiota remains considerably less studied. Crucially, as in mammals, microbial communities of the avian gut have been linked to host health, fitness, and behaviour, representing both key biological processes and central research themes in ecology to conservation^4,5,10,21–23^.

Early life represents a critical window for microbiota establishment, with lasting consequences for host development and health^24,25^. For the majority of mammals, microbial seeding begins at birth when the previously sterile gut is colonised during passage through the birth canal^26–28^, and continues via maternal transfer (largely through lactation) and environmental acquisition^29,30^. These tightly regulated exposure routes are thought to contribute to broadly conserved age-related microbiota trajectories across several mammalian species, including increasing richness and evenness during early development^12,14,17,31–33^ (but see Reese et al., 2021^34^). Birds, however, follow fundamentally different patterns due to their distinct reproductive biology and early-life environments. Birds are oviparous and lack these maternal microbiota transmission routes. Instead, early-life microbial acquisition is likely shaped by the environment (e.g. soil, nest materials) and social interactions such as feeding (parent to offspring). These more variable transmission routes may lead to less predictable microbiota trajectories during early development. This hypothesis is supported by studies showing variable age-related changes in alpha diversity across bird species^21–23,35,36^ (for comparison: increasing richness and evenness during early development is observed in several mammalian species^14,17,29,31–33^ (but see Reese et al., 2021^37^)). These fundamental differences in reproductive biology mean that mammalian findings cannot be directly extrapolated to birds, positioning avian species as essential comparative models for understanding microbiota assembly across vertebrates.

In this study, we investigate the gut microbiota of the Atlantic puffin (*Fratercula arctica*), a long-lived, migratory seabird with ecology that makes it particularly suited for studying how age and environment shape the gut microbiota beyond mammalian hosts. Specifically, chicks develop in underground burrows, where they likely acquire microbes from soil and nest environments. They are also provisioned with a relatively narrow range of marine prey items carried in their parents’ beaks^38^, and are thus dosed with microbes from both parents and their diet. In contrast, adults forage widely at sea and migrate between breeding seasons, potentially encountering common environmental microbial sources across populations^39^. Puffins are also of particular research interest as they are increasingly vulnerable to environmental change, with population declines across their North Atlantic range leading to their IUCN listing as ‘vulnerable’^20^ and inclusion on BirdLife International’s European Red List of Birds (2021)^40^. Furthermore, their overlapping diet and breeding habitats with several other North Atlantic seabird species make insights from puffin microbiota potentially applicable to understanding gut microbial ecology across a broader range of seabirds. We used 16S rRNA amplicon sequencing of faecal samples collected from chicks and adults across six British populations spanning ecologically distinct marine areas (Celtic Sea, Hebrides, Orkney, Shetland, Firth of Forth). We examined how age and geography influence gut microbiota diversity, composition, and aerotolerance as measures of microbiota structure and function.

## Methods

### Sample collection

A total of 153 faecal samples were collected in July 2021 from seven British populations; Skomer Island (Pembrokeshire, Wales), Skokholm Island (Pembrokeshire, Wales), Fair Isle (Shetland Islands, Scotland), Shiant Isles (Hebrides, Scotland), Sule Skerry (Orkney Islands, Scotland), Pentland Skerries (Orkney Islands, Scotland), and Isle of May (Firth of Forth, Scotland) (Supplementary Table 1; see map of islands for which samples were successfully sequenced in Figure 1B). Samples were collected from adults and chicks. Chicks were estimated to be around 2–5 weeks old at the time of sampling, still residing in their burrows and dependent on parental provisioning. Age could not be determined for adults beyond their status as breeding individuals, and sex could not be determined for either chicks or adults. Samples were collected opportunistically during routine monitoring of nests or annual ringing campaigns by briefly holding the animal over a sterilised sampling dish. All sample collection was conducted by field personnel licenced for bird handling under UK regulations. Faecal samples were collected from a sterile sampling dish using a sterilised spatula. Samples were preserved in DNA/RNA Shield (Zymo Research, USA), shipped to the laboratory at ambient temperature, and stored at -80°C until DNA extraction (5 months).

**Figure 1.**
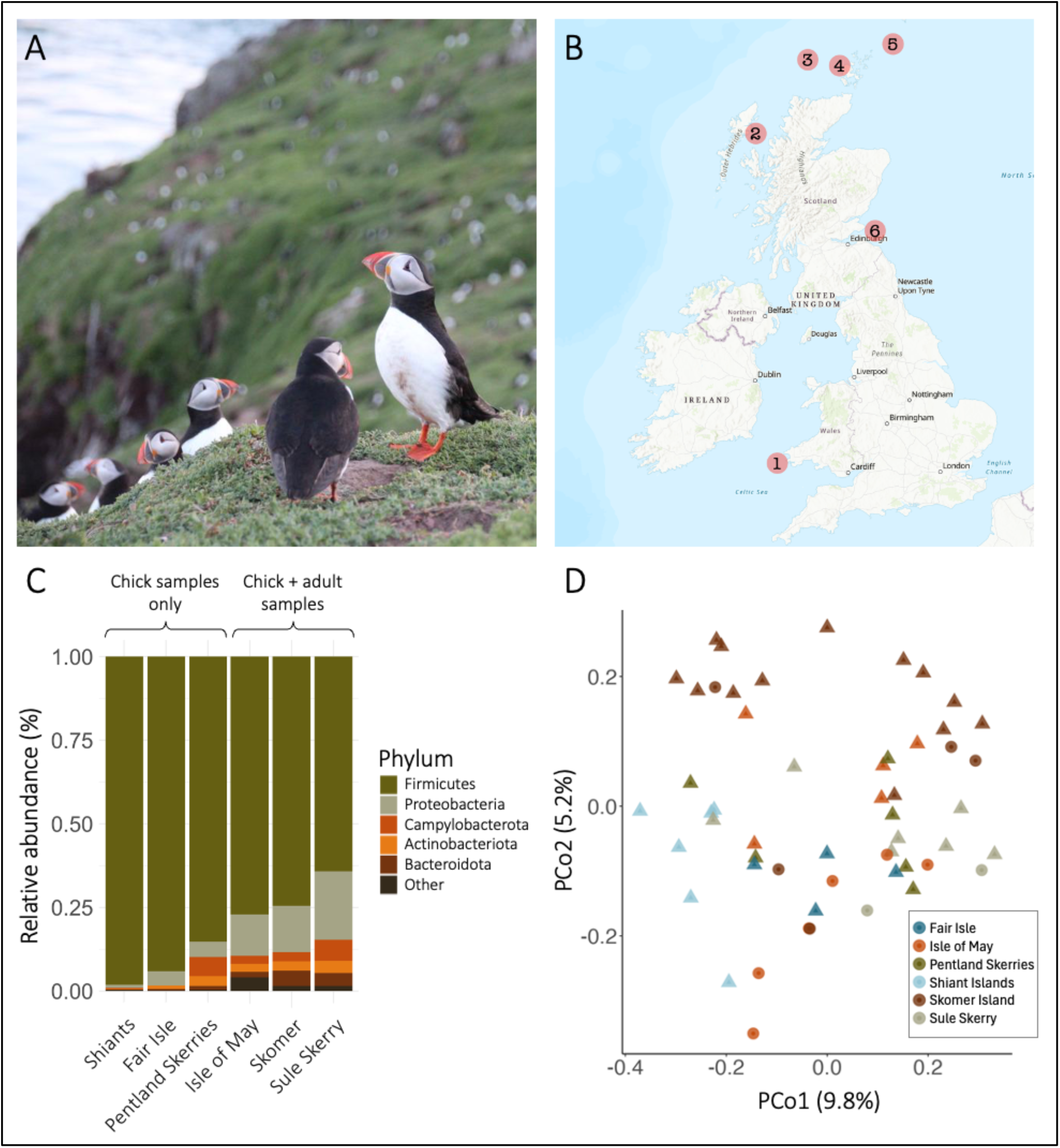
(**A**) Atlantic puffin (*Fratercula arctica*) (photo: Eveliina Hanski). (**B**) Map showing geographic locations of six British Atlantic puffin populations included in the study: 1. Skomer Island; 2. Shiant Isles; 3. Sule Skerry; 4. Pentland Skerries; 5. Fair Isle; 6. Isle of May. (**C**) Mean relative abundance of bacterial phyla across six populations, ordered by decreasing relative abundance of Firmicutes, the dominant phylum. Rare taxa (mean relative abundance <5% and prevalence <5% across samples) are under ‘Other’. (**D**) Principal coordinate analysis of adult (*n*=13) and chick (*n=*41) puffins on Jaccard distance. Microbial community similarity between sample pairs increases with point proximity. Shape indicates age category; *circle =* adult, *triangle =* chick.

### Laboratory work

Faecal samples were randomised into eight batches of up to 23 samples for DNA extraction. DNA was extracted using ZymoBIOMICS DNA MiniPrep kit (spin-column format) according to manufacturer’s instructions (Zymo Research, USA). A negative control (DNAse-free H2O) was included in each DNA extraction batch at a varying position to account for potential systematic cross-contamination between positions. Microbiota profiling was conducted using amplicon sequencing, targeting the V4–V5 region of the 16S rRNA gene using primers 515F and 926R^41,42^. Library preparation was conducted in two 96-well plates of up to 94 samples (comprising true samples as well as negative controls). Each library preparation also included a negative PCR control (at a varying position). Amplicons were sequenced in a single sequencing run using the Illumina MiSeq platform (Reagent kit v3, 2x300 bp chemistry). Library preparation and sequencing was conducted at the Integrated Microbiome Resource, using the protocol described in Comeau et al. (2017)^43^. All negative controls for DNA extraction and PCR reactions were sequenced intermixed with true samples, alongside the sequencing facility’s negative reaction for the amplicon sequencing.

### Data processing

Data processing and subsequent analysis was conducted using R v4.1.2. Demultiplexed sequencing reads were processed through the *DADA2* pipeline^44^. Taxonomy was assigned using the SILVA rRNA database v138.1^45^. Packages *DECIPHER*^46^ and *phangorn*^47^ were used to build a phylogenetic tree. A total of 8 unique ASVs were detected across all negative DNA extraction and library preparation controls (*n*=9), with a maximum read count of 77 for any given control (range 0–77, mean 15; for comparison, mean read depth was 66,176 for true samples). Using these negative controls, we tested for the presence of any potential contaminants using the R package *decontam*^48^ using the `prevalence` method, in which the prevalence of each sequence in biological samples is compared to that in negative controls (conducted separately with extraction and library preparation controls). A sequence was considered a contaminant if it reached a probability of 0.1 in the Fisher’s exact test used in *decontam*. Across the two tests (decontam with either extraction or library preparation controls), one ASV contaminant was identified and filtered from the data before subsequent analysis.

The package *iNEXT*^49^ was used to generate sample completeness and rarefaction curves, which were used to determine a threshold for sufficient sequencing depth. With this, the sequencing depth threshold was set to 5,000 and samples under the threshold (*n*=99) were consequently removed from the data. Asymptotic ASV richness and Shannon diversity estimates were also measured with *iNEXT*, after which ASVs that were never observed at a count greater than 1 in any individual sample were removed from the dataset (*n*=245). The package *phyloseq*^50^ was used to normalise ASV counts to proportional abundance. Finally, two duplicates (two faecal depositions were sampled twice) were removed from the data. After data processing, 54 unique samples from six populations were retained for analysis (Table S1) with a mean read depth of 66,176 (range 5,447–198,796). Across these samples, 2,096 unique amplicon sequence variants (ASVs) were detected, with each sample having 113 unique ASVs on average (range 11–277). Both adult and chick samples were available for analysis in three of the six populations (Skomer, Isle of May, and Sule Skerry; Table S1).

### Phenotypic profiles of bacteria

Bacterial aerotolerance was determined using a custom database as described in Hanski et al. (2024)^17^. Briefly, bacterial genera were categorised into ‘obligate anaerobes’ (when specifically listed as *obligate* anaerobes) and ‘aerotolerants’ (for all facultative anaerobes, microaerobes, facultative aerobes and aerobes) or ‘unknown’ (when information was not available) using Bergey’s Manual of Systematics of Archaea and Bacterial or additional peer-reviewed references when information was not available in the manual^51^. The aerotolerance data used here is available online (see *Data availability*).

### Data analysis

To investigate the proportion of gut microbiota variation explained by population and age (adult vs chick), we performed marginal permutational multivariate analyses of variance (PERMANOVA) on Jaccard and Aitchison distances using the `adonis2` function in R package *vegan*^52^. Jaccard distance was calculated using the `distance` function in package *phyloseq,* with the `binary` argument set to `TRUE`, such that relative abundances of taxa are not considered. Aitchison distance was calculated by first applying a centered log-ratio (CLR) transformation using the `transform` function from the package *microbiome*^53^. This CLR transformation replaces exact zero relative abundances with a pseudocount (min(relative abundance)/2) before taking logarithms. The `distance` function from the package *phyloseq* was then used with the method set to `euclidian`. DNA extraction batch and read depth were included as predictors. Beta dispersion by population ID and age category was tested for using the `betadisper` function in package *vegan*^52^, a method that is sensitive to unbalanced group sizes.

To study predictors of gut microbial similarity between sample pairs, we used dyadic Bayesian regression framework (function `brm`) with R package *brms*^54^. It comprises a pairwise generalised linear mixed model predicting gut microbial similarity (on Jaccard distance, measured as described above) with other measures of a sample pair (e.g., spatial proximity). We ensured that similarity scores (Jaccard indices) fell strictly within the (0,1) interval by applying a standard transformation: (x × (n − 1) + 0.5) / n, where n is the sample size. We then used logit link functions of family *beta*. The effect of spatial proximity on microbiota similarity was estimated separately for (1) *adult–adult* and (2) *chick–chick* sample pairs with two models. As the sample size for adults was substantially smaller than that of chicks (Table S1), we repeated the chick model using a subset of randomly selected sample pairs matched in population representation and sample size to the adult dataset, with the only exception that Isle of May had five instead of six samples (Table S1). Models used population ID similarity (same vs different population) and at-sea proximity (puffins do not fly over land, so at-sea distance represents a more ecologically relevant distance between populations than absolute distance) as predictors. For this, at-sea distance was scaled to the [0,1] range as described above and converted to proximity (1 – normalised distance) for more intuitive interpretation. These spatial analyses were repeated on the anaerobic and aerobic communities of the microbiota (as aerobic taxa predominantly spread through environmental exposure while anaerobic taxa spread through social transmission^55^), with difference in the relative abundance of taxa with unknown aerotolerance included as a fixed effect.

For investigation of microbiota variation across age categories, we used two approaches. First, we predicted microbial similarity with pairwise age category similarity (same vs different age category). Second, to estimate whether beta diversity varies across the two age categories, we predicted microbial similarity with pairwise age category (*adult–adult* or *chick–chick*; only sample pairs with same age category were included). Both models investigating age effects on microbial similarity included population ID similarity as a covariate. For identifying taxa driving age-effects in the microbiota, we used a ‘drop-one-taxon-out’ approach. Here, ASVs associated with a single genus were excluded before re-measuring the Jaccard index, followed by running the brm model as described above (`Jaccard distance ∼ pairwise age category + …`). This was repeated until each genus and its associated ASVs had been dropped out from the data at a time. We then quantified ‘importance’ scores for each of the 465 genera, as described in Raulo et al. (2024)^18^, reflecting the extent to which excluding a genus from the data reduced the certainty of the age effect estimate (i.e., increased the credible interval width).

All dyadic beta regression models included DNA extraction batch similarity (same vs different extraction batch) as well as read depth difference of the sample pair as fixed effects. To account for the inherent autocorrelation of pairwise values, all dyadic models included a multi-membership random effect `(1|mm(ID1, ID2)`^18^.

To assess whether alpha diversity varied across age categories, we fitted two separate models for asymptotic Shannon diversity and asymptotic ASV richness (estimated with *iNEXT*^49^, see details in *Data processing*) using *brms*^54^. For Shannon diversity, we used a *zero-inflated beta* regression model, with a precision (phi) term predicted by age to account for potential differences in dispersion. The response variable was scaled to the [0,1] range. For ASV richness, we fitted a *Gaussian regression model* with a log-transformed response. In both models, age category (adult vs chick) and read count were included as fixed effects, with population ID as a random effect to account for population-level variation. Alpha diversity models were run with (1) all samples and (2) all except Shiants samples, since observed ASV richness was substantially lower in Shiants population (Table S1).

To examine whether microbiota aerotolerance varied by age (anaerobe:aerobe ratios have been observed to increase with age in several mammalian species^14,17,56^), we similarly implemented a brm model. The proportion of aerotolerant taxa out of taxa with known aerotolerance was used as the response variable. Age category, scaled read count, DNA extraction batch, and summed relative abundance of taxa with unknown aerotolerance were included as fixed effects, while population ID was included as a random effect to account for variation across populations. Read count was scaled to improve model convergence and ensure comparability across samples. Since the response variable included values of exactly 1, we specified a *zero-inflated beta* distribution to appropriately model the data.

For all brm models, posterior checks were conducted to confirm reliable model performances. Here, we ensured that the chains had converged, Rhat values were <1.05, bulk effective sample sizes at least 100 times the number of chains and no smaller than 10% of total posterior draws, and that the sampler took small enough steps to avoid excess (>10) divergent transitions after warm-up. Additionally, posterior predictive checks were performed to assess model fit, confirming that the simulated posterior distributions closely matched the observed data, with only minor deviations in certain regions.

ASV networks were constructed to visualise microbiota similarity between puffin populations based on chick samples. Only chick samples were included, as adult samples were available for only three of the six populations (Table S1). Networks were generated separately for the whole microbiota, anaerobic taxa, and aerobic taxa (aerotolerance determined at genus level, see above). To ensure comparability across populations with uneven sample sizes, networks were based on 100 iterations of random subsampling of four chick samples per population (the sample size available in the population with the fewest chick samples; Table S1). In each iteration, presence–absence data were used to calculate ASV richness within populations and Jaccard similarity (*n* ASVs present in both populations / total *n* ASVs across the two populations) between population pairs. Final node sizes and edge thicknesses represent the average ASV richness and Jaccard index, respectively, across all iterations.

## Results

### Atlantic puffin gut microbiota composition and structure across six British populations

We profiled the bacterial component of the Atlantic puffin gut microbiota in adults (*n*=13) and chicks (*n*=41) from six populations in the UK. At the phylum level, the puffin gut microbiota was dominated by Firmicutes (mean relative abundance 78.6%) and Proteobacteria (11.5%), followed by Campylobacterota (3.1%), Bacteroidota (2.4%) and Actinobacteriota (2.4%) (Fig. 1C, Fig. S1A). The most common genera were *Catellicoccus* (43.1%), *Clostridium sensu stricto 1* (22.2%)*, Escherichia-Shigella* (5.3%), and *Paeniclostridium* (5.0%) (Fig. S1A, Fig. S2B).

Both population and age category (adult vs chick) significantly explained variation in microbiota composition, though age had markedly lower explanatory power than population. Population explained 15.6% of variation in the taxa detected in the samples (marginal PERMANOVA; binary Jaccard index, *population*: *R^2^*=0.156, *F*=1.838; *p*=0.001, *age*: *R^2^*=0.023, *F*=1.337; *p*=0.032; beta dispersion, *population*: *F*=4.511, *p*=0.005, *age*: *F*=2.997, *p*=0.084; Fig. 1D) and 13.6% of variation in the relative abundances of these taxa (Aitchison distance, *population*: *R^2^*=0.136, *F*=1.661; *p*=0.001, *age*: *R^2^*=0.030, *F*=1.788; *p*=0.008; beta dispersion, *population*: *F*=2.223, *p*=0.073, *age*: *F*=1.766, *p*=0.191; Fig. S2B).

Overall, the puffin gut microbiota was dominated by aerotolerant taxa (mean relative abundance 70.4%, sd 29.3%; mean relative abundance of anaerobic taxa 31.5%, sd 34.0%; Fig. S1C, Fig. S2C; aerotolerance was determined at genus level, see Methods).

### Age-related variation in microbial diversity and composition

Microbiota composition differed significantly between adults and chicks. Sample pairs from the same age category were more similar than those from different age categories (Bayesian regression model (brm): posterior mean for *same* = 0.12, 95% credible intervals (CIs) 0.02 to 0.21; see details for all models in Methods). Among same-age pairs, chick microbiotas were more similar to one another than adult microbiotas (posterior mean for *chick-chick* pairs = 0.94, CIs 0.50 to 1.38), suggesting beta diversity increased with age (Fig. 2A; see also individual-level phylum and genus profiles in Fig. S1).

**Figure 2.**
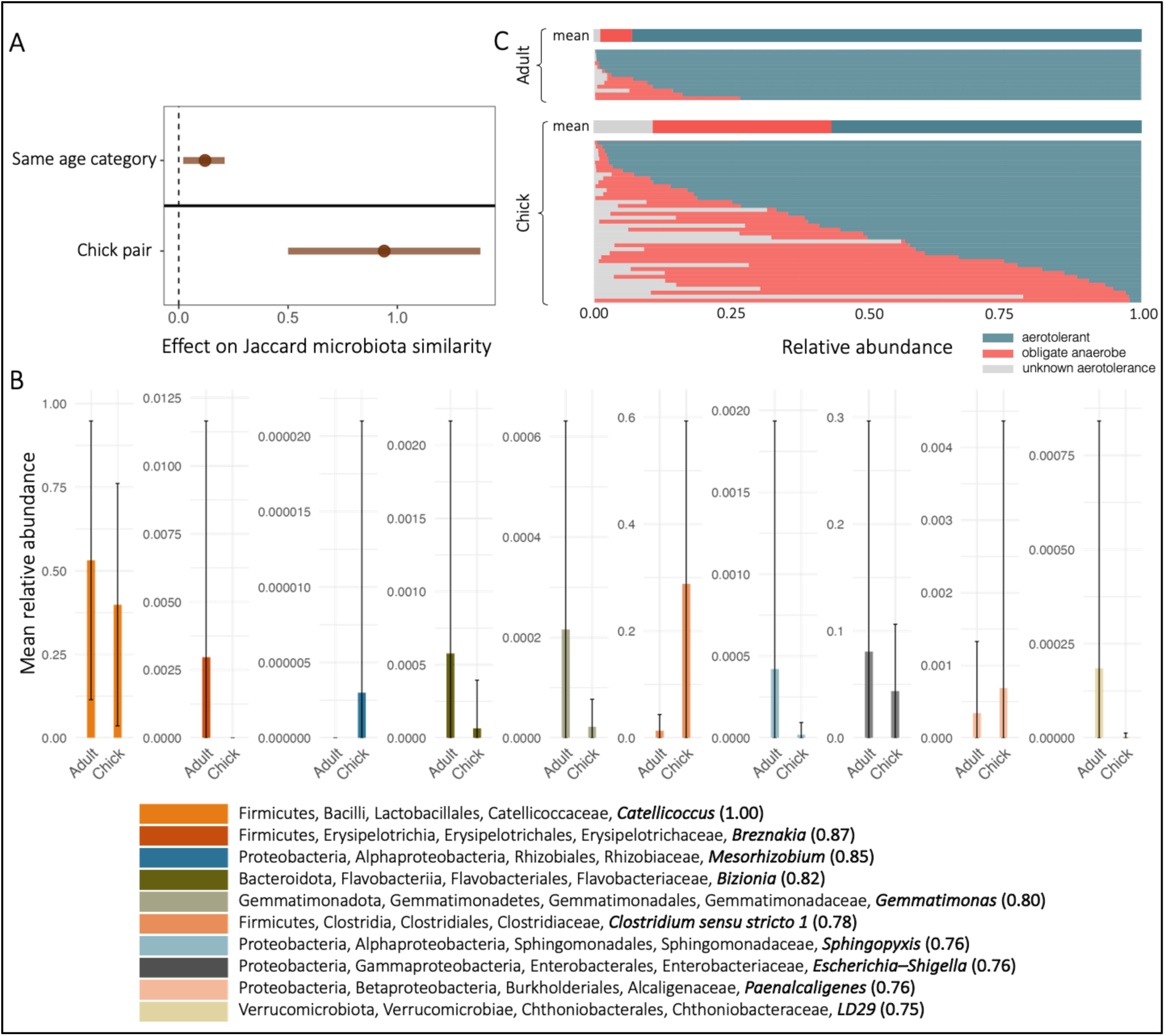
(**A**) Estimated effects of age similarity (*top*) and category (*bottom*) on pairwise gut microbiota similarity (Jaccard distance), based on two Bayesian regression models (brm). The top model (“same age category”) estimates the effect of whether two samples are from the same vs different age categories (reference: different age). The bottom model (“chick pair”) estimates the effect of being a chick pair vs an adult pair, among within-age comparisons (reference: adult pair). Points show posterior means; horizontal lines show 95% credible intervals (CIs). A variable significantly predicts community similarity if its CI does not overlap zero. The effect sizes of the estimates between the two models are not directly comparable. Y-axis labels (‘Same age category’, ‘Chick pair’) indicate the predictor levels being compared to their respective reference groups. (**B**) Mean relative abundance (± standard deviation) of the top 10 bacterial genera contributing most strongly to age-related differences in microbiota composition, identified using a ‘drop-one-taxon-out’ approach (see Methods). Importance scores (in parentheses) quantify the influence of each genus on the age effect estimate. (**C**) Stacked bar plots showing the aerotolerance profile of gut microbiota across individual puffins, grouped by age. Each thin bar represents an individual, ordered by the relative abundance of aerotolerant taxa within age category. The wider bars at the top of each cluster show the mean relative abundances for adults (top) and chicks (middle). Colours indicate bacterial oxygen tolerance category: aerotolerant (*blue*), obligate anaerobe (*red*), and unknown aerotolerance (*grey*).

In terms of alpha diversity, Shannon diversity was significantly lower in chicks compared to adults (posterior mean for *Chick* = -1.99, CIs -2.85 to -1.13), and also more consistent (i.e., lower variability across individuals; posterior mean *phi* = 4.06, CIs 3.27 to 4.86). However, ASV richness did not differ significantly between age groups (posterior mean for *chick* = 0.03, CIs -0.10 to 0.17). To assess whether the relatively low ASV richness in the Shiants population (mean = 36.9, compared to 95–145 in other populations; Table S1) may have influenced these patterns, we repeated the models excluding this population. Results were nearly identical: Shannon diversity remained significantly lower in chicks (posterior mean for *chick* = -1.94, CIs -2.80 to -1.08), while ASV richness still showed no significant age-related difference (posterior mean for *chick* = 0.06, CIs -0.08 to 0.19).

To further characterise age-related differences in the puffin microbiota, we next examined taxonomic composition. Overall, the relative abundance of Proteobacterial taxa was higher in adults than in chicks (Fig. S1A), while at the genus level, chicks had higher abundances of *Clostridium sensu stricto 1*, which was also among the top 10 genera with the highest importance scores in driving the age effect on the puffin microbiota (Fig. 2B, Table S2; Fig. S2B). These genera were taxonomically diverse, spanning five distinct phyla: Bacteroidota, Firmicutes, Proteobacteria, Gemmatimonadota, and Verrucomicrobiota. This approach assesses how much each taxon contributes to age-related microbiota differences by systematically removing them from the data and measuring the change in model certainty. While several of these taxa had low mean relative abundances (<0.2%) in both adult and chick samples, others, such as *Catellicoccus* (∼53% in adults, ∼40% in chicks) and *Escherichia– Shigella* (∼8% in adults, ∼4% in chicks), were substantially more abundant on average, although their relative abundances varied widely across individuals within each age category (Fig. 2B, Table S2; Fig. S1B). This compositional divergence extended to microbial oxygen tolerance: chicks had a significantly lower proportion of aerotolerant bacteria than adults (posterior mean for *chick* = -1.81, CIs -2.79 to -0.81; Fig. 2C, Fig. S1C, Fig. S2C).

### Puffin chicks but not adults show spatially structured gut microbiota shaped by oxygen tolerance

Spatial effects on microbiota similarity were strongly age dependent. In adults, neither population similarity (same vs different population) nor at-sea proximity significantly predicted microbiota similarity (posterior mean for *same population* = 0.43, CIs -0.44 to 1.30; posterior mean for *at-sea proximity* = -0.38, CIs -1.30 to 0.53; see at-sea population distances in Fig. S3). In chicks, however, microbiota similarity was significantly higher in sample pairs from the same rather than different populations while at-sea proximity had no significant effect (posterior mean for *same population* = 0.79, CIs 0.62 to 0.96; posterior mean for *at-sea proximity* = -0.08, CIs -0.28 to 0.13). These effects were maintained in a reduced dataset matched to the adult sample structure, though the population effect was weaker (posterior mean for *same population* = 0.60, CIs 0.03 to 1.19; posterior mean for *at-sea proximity* = -0.20, Cis -0.81 to 0.41; Table S1).

When splitting the microbiota into anaerobic and aerobic components, additional spatial patterns emerged. In chicks, population similarity and spatial proximity both predicted anaerobic microbiota similarity (posterior mean for *same population* = 0.20, CIs 0.04 to 0.36; posterior mean for *at-sea proximity* = 0.36, CIs 0.16 to 0.55), while aerobic microbiota were more similar within populations but became less similar between populations that were geographically closer (posterior mean for *same population* = 0.76, CIs 0.58 to 0.93; posterior mean for *at-sea proximity* = -0.22, CIs -0.42 to -0.01; Fig. 3A). Although this latter effect was only marginally significant, it suggests both microbial groups show spatial structuring beyond population similarity. However, these opposing spatial patterns may effectively cancel each other out in whole-community analyses, obscuring broader spatial trends (Fig. 3A). In adults, by contrast, neither anaerobic nor aerobic microbiota showed significant spatial structuring (Fig. 3A).

**Figure 3.**
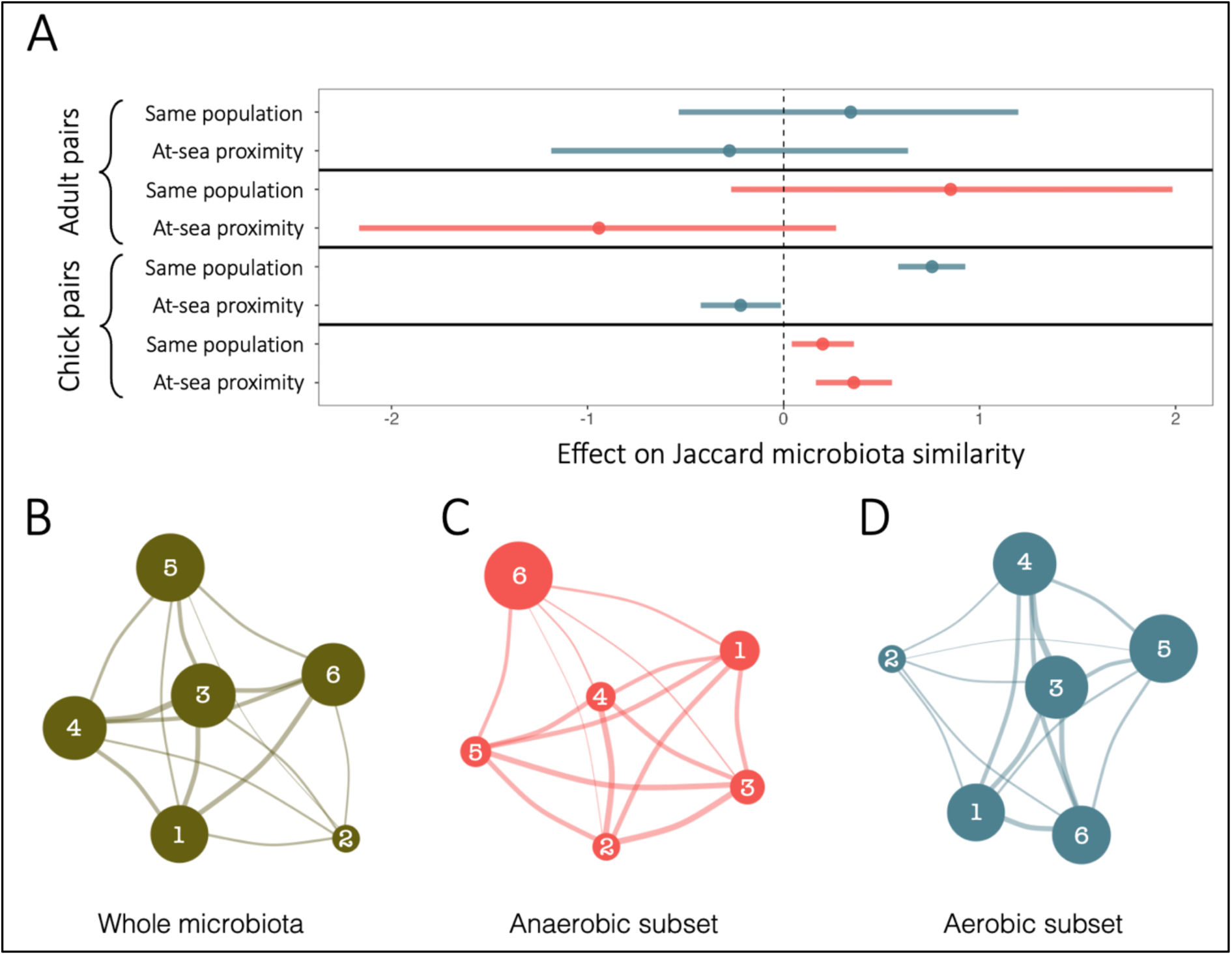
(**A**) Estimated spatial effects on pairwise gut microbial similarity on Jaccard distance in adult and chick sample pairs from Bayesian regression models (brm) that predicted pairwise similarity of aerobic (*blue*) or anaerobic (*red*) microbiota communities in adult-adult or chick-chick sample pairs with population ID similarity (same vs different population with ‘different population’ as reference category) and at-sea proximity (scaled to 0–1 for easier interpretation). Points are posterior means and lines are 95% credible intervals (CIs). A variable significantly predicts community similarity if its credible interval does not overlap zero, and effects are significantly different if their credible intervals do not overlap. Black horizontal lines separate different models. Effect sizes are model-specific and not directly comparable across models. (**B–D**) ASV sharing networks between chick samples from puffin populations (1. Skomer; 2. Shiant Isles; 3. Sule Skerry; 4. Pentland Skerries; 5. Fair Isle; 6. Isle of May) for the whole microbiota (**B**), anaerobic microbiota (**C**), and aerobic microbiota (**D**). Nodes represent puffin populations, with node size proportional to the average ASV richness per population based on 100 iterations of randomly sampled sets of four chick samples. Edges represent pairwise microbiota similarity between populations, quantified as the mean Jaccard index across the same iterations. Edge thickness is scaled to average Jaccard index values. Edge length is determined by a force-directed layout and reflects overall network structure, but does not correspond to spatial distance or microbiota similarity.

To better understand microbiota differentiation between puffin populations, we visualised ASV networks to identify common taxa across individuals. Networks were constructed for the whole microbiota (Fig. 3B) as well as for anaerobic and aerobic communities separately (Figs. 3C and 3D, respectively). The full network revealed strong ASV connectivity among most populations, but also highlighted reduced overlap with the Shiant Isles (node 2), suggesting some population-specific distinctiveness. When partitioned by oxygen tolerance, these patterns diverged. Anaerobic microbiota exhibited broadly connected and relatively even distribution across populations (indicated by the relatively uniform edge thicknesses between nodes). In contrast, the aerobic microbiota exhibited a pronounced separation of the Shiant Isles, with fewer common ASVs with other populations. This suggests that the observed negative effect of spatial proximity on aerobic microbiota similarity (Fig. 3A) may be largely driven by this single, distinct population rather than a consistent pattern across all populations.

## Discussion

Birds are key contributors to ecosystem function^57^, yet many of the host traits supporting these benefits are tightly linked to the gut microbiome^58^ – a component of avian biology that remains underexplored. Using samples from six Atlantic puffin populations from the British Isles, we demonstrate that microbiota composition and structuring is strongly age dependent and displays patterns that suggest divergence from mammalian models.

Overall, the puffin gut microbiota was dominated by Firmicutes and Proteobacteria, with relatively low levels of Bacteroidota. This composition resembles that of other seabirds such as shearwaters^4^, fulmars^4^ and thick-billed murres^5^, as well as Seychelles’ warblers^10^ and bats^59,60^, and is broadly consistent with prior observations from UK puffins that found Proteobacteria and Firmicutes among the dominant phyla^61^. In contrast, gut communities in nonvolant terrestrial mammals are typically dominated by Firmicutes and Bacteroidota^14,17,62,63^, while in marine mammals show mixed patterns^64–67^, suggesting that marine ecology alone does not consistently drive high Proteobacteria abundance.

Age-related microbiota dynamics in birds are of particular interest due to their distinct developmental context, likely shaped by environmental exposure rather than the primary maternal transmission routes (birth canal and lactation) that influence mammalian microbiota^14,17,68,69^. This may result in more variable early-life microbial dynamics across individuals, populations, and bird species, in contrast to the relatively conserved trajectories seen in several mammalian species^14,17,70^. Our findings support this view, revealing clear age-related shifts in gut microbiota diversity and composition.

Shannon diversity was significantly higher in adults than in chicks (who were ∼2–5 weeks old at sampling), while ASV richness remained comparable, suggesting that evenness, rather than richness, increases with age. This pattern diverges from mammals, where both richness and evenness typically increase during early development^12–14,17^, but aligns with more heterogenous age dynamics reported in other bird species^21,22,35,36,71,72^. The increased evenness (reflected in higher Shannon diversity) in adults may reflect dietary diversification and immune system maturation, which could promote more even microbial communities. One possible explanation for the lack of richness differences is that since chick samples were collected weeks after hatching, initial colonisation dynamics may have already been stabilised, limiting our ability to capture early richness changes.

These age-related diversity patterns were accompanied by marked compositional differences between chicks and adults, driven primarily by *Catellicoccus*, which was the most abundant genus in our dataset and showed significantly higher relative abundances in adults. The dominance of *Catellicoccus* in puffins has been reported previously^61^ and is also observed in gulls^73^ and thick billed murres^5^, with detected in various waterfowl^74,75^. The only described species, *Catellicoccus marimammalium,* exhibits features of a host-adapted lifestyle, including limited biosynthetic capacity and specialised nutrient transport systems^76^, and has been associated with longer telomere length in yellow-legged gulls^77^, hinting at potential links to host health. Originally isolated from porpoise and grey seal^78^, *Catellicoccus* has been primarily reported in seabirds and marine mammals, and appears to be uncommon in terrestrial mammals. This distribution pattern suggests that shared habitat (rather than host phylogeny alone) may influence its prevalence in marine-associated mammals. In puffins, its higher relative abundance in adults could reflect increased marine exposure. Given its dominance in multiple avian taxa, signs of host adaptation, and age-association across species, *Catellicoccus* may represent a ‘core’ member of the avian gut microbiota with functional importance.

Interestingly, we found that the overall composition of gut microbiota became increasingly individualised with age. Chick microbiotas were more similar to one another than those of adults, suggesting that microbial communities diverge between individuals as they grow older. This pattern is consistent with findings in some other bird species^22^ but contrasts with patterns observed in common mammalian systems (primates and mice), where early-life microbiotas tend to be more variable between individuals and converge over time^13,14,17^. The greater similarity in chick microbiotas may reflect their narrow age range and more constrained environments compared to adults: chicks were all sampled within a narrow age range, whereas adults likely spanned a much wider age range (puffin breeding age can range ∼5-40 years). Furthermore, adults also forage widely^79,80^ and migrate across large marine areas throughout the year, with overlap between the non-breeding distribution of different populations but substantial individual differences^39^, potentially encountering similar environmental microbes regardless of their breeding population, which may reduce population-level differences while still increasing variation between individuals.

Indeed, spatial structuring of the microbiota was evident in chicks but not adults. Chick microbiotas were more similar within than between populations, consistent with local environmental shaping, while adult microbiotas showed no such geographic structure. This age-dependent spatial pattern supports the hypothesis that avian microbiota assembly is more environmentally driven than in mammalian species. Puffin chicks remain in underground burrows for several weeks post-hatching^81^ and are therefore exposed to locally distinct soil microbial communities. These early-life exposures likely shape the initial microbial community, contributing to the population-level signal observed. In early life, microbial communities are also thought to be under weaker host physiological control^82–84^, making them more susceptible to local environmental influences. As hosts mature, increased immune modulation and physiological filtering may contribute to greater host-driven divergence, as well as a weakening of geographic signals. In puffins, this may help explain why adult microbiotas are both more variable across individuals (i.e., higher beta diversity) and less spatially structured. The absence of spatial structure in adults may also partly reflect our smaller adult sample size, though when chick analyses were restricted to match the adult sample structure, a population effect was still marginally detectable.

To further understand the ecological drivers shaping puffin gut microbial communities, we partitioned the microbiota by oxygen tolerance. Chicks harboured a higher proportion of obligate anaerobes, whereas adults were dominated by aerotolerant taxa. This contrasts patterns observed in mammals^17,69,82^ and may be explained by a combination of ecological, physiological, and/or behavioural factors unique to puffins or potentially seabirds more broadly. For instance, adult puffins spend significant time foraging at or near the ocean surface, where they are exposed to bacteria that are likely aerotolerant due to the high oxygen levels in surface seawater. While chicks are similarly exposed to soil microbes, which are likely to be aerobic, their exposure may well be more passive, as they are unlikely to actively ingest soil in the way adults may take in seawater while feeding. Further research is needed to determine whether the observed pattern in the ratio of anaerobes to aerobes is common in seabirds and what factors (e.g., immune modulation, feeding frequency) might be driving it. It is also important to note that in our cross-sectional microbiota profiles we cannot distinguish whether detected microbes – including the aerotolerant taxa in adults – represent resident members of the gut microbiota or transient taxa associated with recent environmental exposures, which could be resolved through longitudinal sampling.

The spatial structuring of the microbiota supports a divergence in how anaerobic and aerobic microbial communities respond to host and environmental factors. In chicks, anaerobic microbiotas were more similar within populations and between geographically proximate populations. In contrast, aerobic microbiota showed higher similarity within populations but decreased similarity with increasing spatial proximity. This negative spatial effect, while only marginally significant, may reflect environmental filtering or localised divergence in aerobes, potentially driven by a single distinct population. The Shiant Isles exhibited highly distinct aerobic microbiota and also had markedly lower ASV richness for both aerobic and anaerobic taxa, suggesting that reduced diversity and unique local conditions – such as island-specific geology or soil microbiota – may have contributed to its differentiation. These findings demonstrate how partitioning microbial communities by functional traits such as oxygen tolerance can reveal ecological structure that remains obscured in whole-community analyses. As oxygen tolerance was classified at the genus level, higher-resolution data will be important to validate these findings and capture potential variation within genera.

Despite yielding considerable advancements in our understanding of the Atlantic Puffin gut microbiota, this study had limitations. Sequencing success was low (65% of samples failed), resulting in small sample sizes with uneven representation of populations and age categories. Unlike mammals, birds excrete nitrogenous waste and gastrointestinal excrement together in the faeces^85^, resulting in low DNA yields and high rates of PCR inhibition^86,87^ – a common challenge for avian microbiota studies^88^. While our cross-sectional comparison of adults and chicks at a single timepoint reveals marked differences between the two age categories, longitudinal sampling from hatching through fledging would be needed to fully capture the microbiota assembly process and initial colonisation dynamics in birds. Future studies would also benefit from investigating microbiota components beyond bacteria, such as fungi (the ‘mycobiota’), as well as exploring high-resolution functional changes in the puffin gut microbiota and their relationships to host health and fitness.

In conclusion, our study provides characterisation of the gut microbiota in the Atlantic puffin, revealing a dynamic interplay between host age, microbiota composition, and spatial ecology. The patterns we document – including population-level structuring in chicks that disappears in adults, increasing individualisation with age, and increased aerotolerance – appear to contrast those observed in commonly used mammalian model systems. These findings highlight the need for further research into the ecological and evolutionary drivers of microbiota variation in birds and suggest age-related microbiota dynamics may be more varied across vertebrates. As avian species face increasing environmental pressures, including global disease threats such as highly pathogenic avian influenza (HPAI), understanding factors driving variation in microbiota composition and what consequences microbiota development has on later health will be increasingly important. More broadly, our results contribute to a growing recognition that birds can serve as valuable comparative models for advancing microbiota research beyond the mammalian species.

## Author information

### Contributions

EH and ALF designed the study. ALF, LS, LS, RH, and MN collected the samples. EH, LT and AR conducted the laboratory work. EH analysed the data. EH, KB, and ALF wrote the manuscript. All authors contributed to the final version of the manuscript.

## Acknowledgements

We thank Rhys Goodhead, Jim Lennon, Stephanie Griffiths, Rebekah Goodwill, Georgia Platt, Dan Gornall, Giselle Eagle, Richard Brown, and Ian Beggs for their help with sample collection. We thank NatureScot for permission to carry out work on Fair Isle and Sule Skerry, the Wildlife Trust for South and West Wales and the Skomer wardens for permission to work on Skomer.

## Data availability

The 16S rRNA amplicon sequencing data used in this study have been deposited in the Sequence Read Archive (SRA) under accession code PRJNA1269590. Sample specific metadata, aerotolerance data, taxonomy table, ASV relative abundance table as well as R code used in the study are publicly available at https://github.com/eveliinahanski/fratercula_arctica.

## Supplementary Material

**Supplementary Figure 1.**
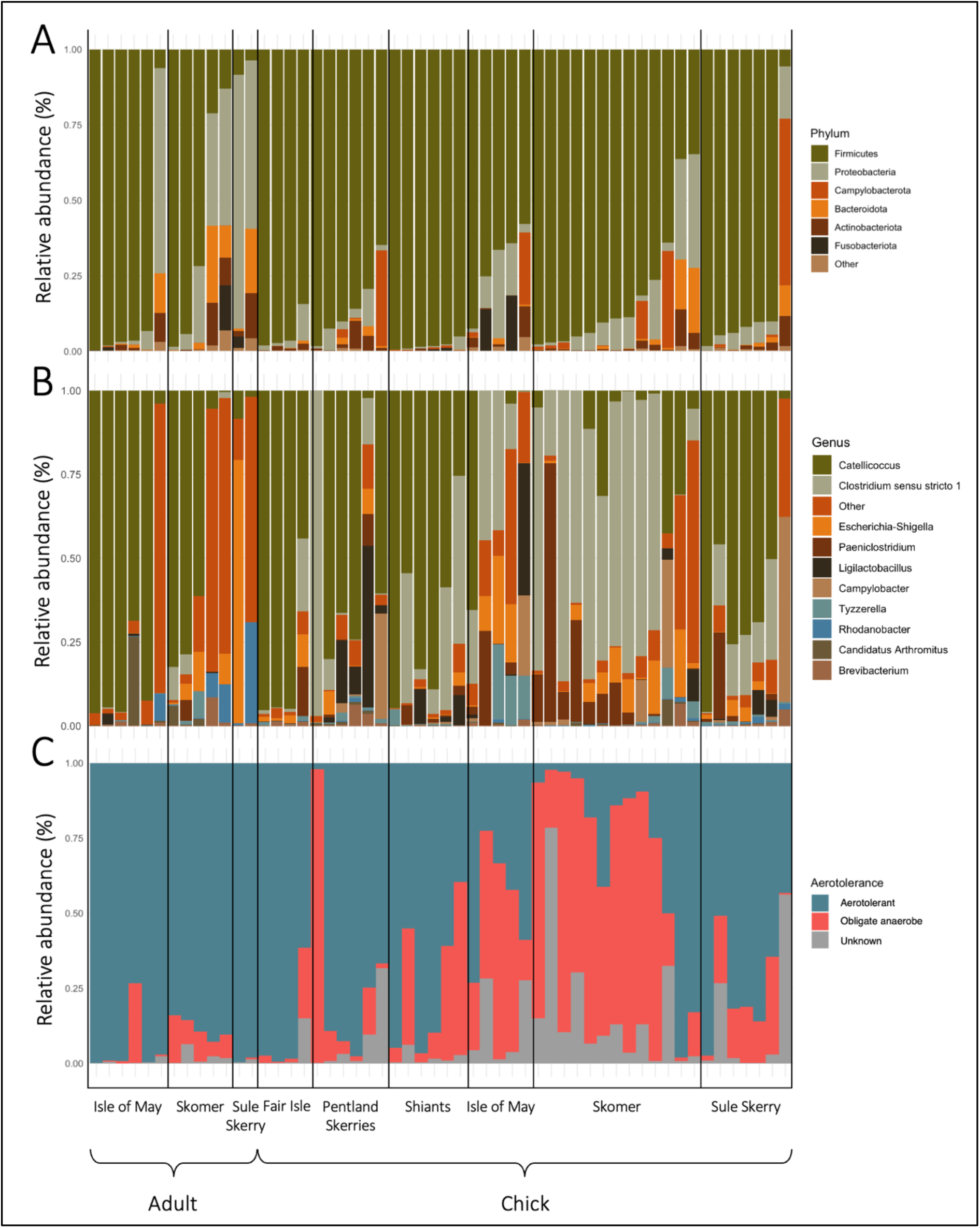
Gut microbiota composition and oxygen tolerance of individual Atlantic puffins. Each vertical bar represents one puffin sample (*n*=54). Samples are grouped by age category (Adult vs Chick) and within age category, ordered by population (Isle of May, Skomer, Sule Skerry, Fair Isle, Pentland Skerries, Shiants). Within each population–age group, samples are ordered by decreasing relative abundance of the dominant phylum, Firmicutes. (**A**) Relative abundance of bacterial phyla across samples. Taxa with mean relative abundance <5% and prevalence <5% are grouped under ‘Other’. (**B**) Relative abundance of bacterial genera across samples. Rare genera (mean relative abundance <5% and prevalence <5%) are grouped under ‘Other’. (**C**) Relative abundance of aerotolerant, obligate anaerobic, and bacteria of unknown aerotolerance across samples. Genera with ambiguous or mixed aerotolerance classifications are grouped as ‘Unknown’.

**Supplementary Figure 2.**
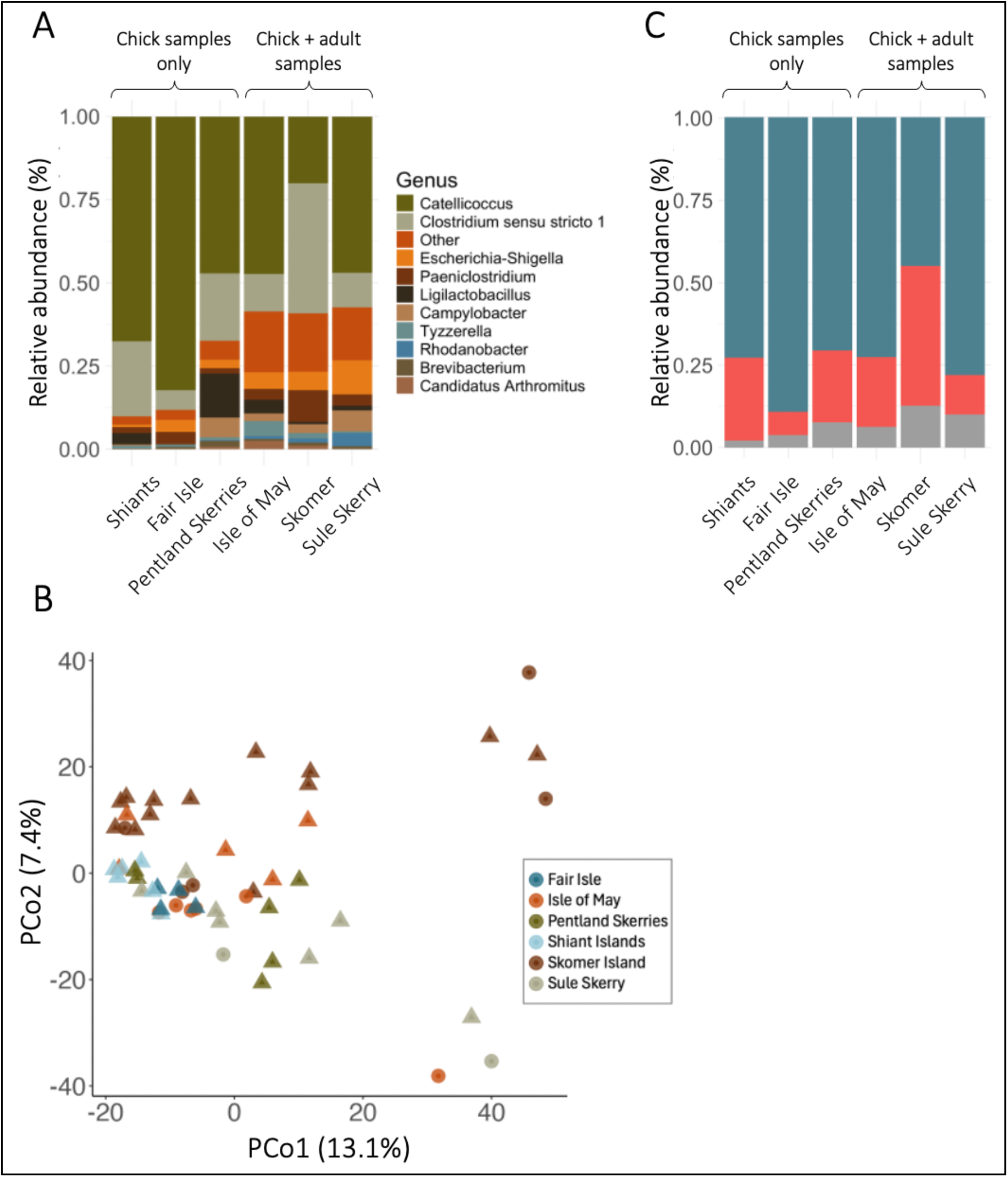
(**A**) Mean relative abundance of bacterial genera across six colonies. Rare taxa (mean relative abundance <5% and prevalence <5% across samples) are under ‘Other’. (**B**) Principal coordinate analysis of adult (*n*=13) and chick (*n=*41) puffins on Aitchison distance. Microbial community similarity between sample pairs increases with point proximity. Shape indicates age category; *circle =* adult, *triangle =* chick. (**C**) Mean relative abundance of aerotolerant and anaerobic bacteria in puffin gut microbiota. Taxa with mixed aerotolerance at genus-level (genera include both aerotolerant and anaerobic bacteria) as well as those with unknown aerotolerance are under ‘Unknown’. Colours indicate bacterial oxygen tolerance category: aerotolerant (*blue*), obligate anaerobe (*red*), and unknown aerotolerance (*grey*).

**Supplementary Figure 3.**
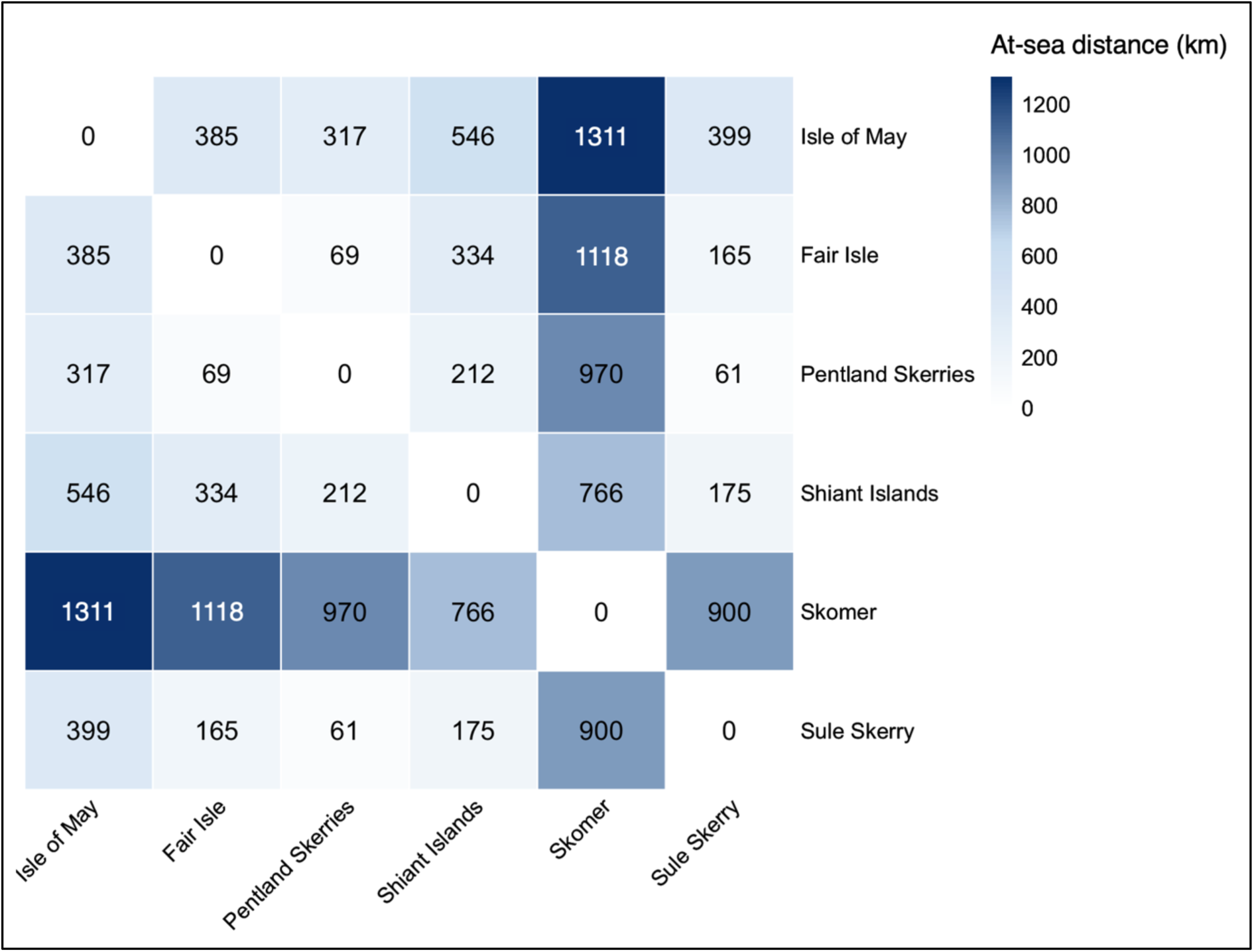
At-sea distances (km) between pairs of Atlantic puffin colonies. Distances represent the shortest continuous travel paths between populations constrained to the marine environment, avoiding land. Colours indicate relative spatial proximity, with darker shades representing greater distances.

**Supplementary Table 1.** Gut microbiota profiles were successfully constructed for 54 samples from six British Atlantic puffin populations. The table reports the number of successfully processed samples for adults and chicks separately, with the number of failed samples shown in brackets. For each group, the range and mean of sequencing read counts are reported, along with asymptotic estimates of ASV richness and Shannon diversity derived from iNEXT (see Methods). These diversity estimates account for variation in sequencing depth and are used in the alpha diversity comparisons reported in the main text.

| Population | Adult |  |  |  | Chick |  |  |  |
| --- | --- | --- | --- | --- | --- | --- | --- | --- |
|  | Nr of<br>successful<br>samples<br>(failed) | Read count<br>range<br>(mean) | Asymptotic<br>ASV<br>richness<br>range<br>(mean) | Asymptotic<br>Shannon<br>diversity range<br>(mean) | Nr of<br>successful<br>samples<br>(failed) | Read count range<br>(mean) | Asymptotic<br>ASV<br>richness<br>range<br>(mean) | Asymptotic<br>Shannon<br>diversity range<br>(mean) |
| *Fair Isle* | 0<br>(0) | N/A | N/A | N/A | 4<br>(11) | 75,370–118,312<br>(101,321) | 47.2–265<br>(128) | 1.52–12.8<br>(4.59) |
| *Isle of May* | 6<br>(1) | 5,491–75,977<br>(46,910) | 11–223<br>(95) | 1.36–134.0<br>(23.9) | 5<br>(5) | 14,270–138,041<br>(60,900) | 60–174<br>(121) | 4.99–20.8<br>(11.7) |
| *Pentland Skerries* | 0<br>(6) | N/A | N/A | N/A | 6<br>(8) | 24,851–142,664<br>(69,666) | 47–276<br>(122) | 3.33–13.8<br>(6.45) |
| *Shiant*s | 0<br>(1) | N/A | N/A | N/A | 6<br>(10) | 7,094–121,209<br>(66,755) | 17–58.1<br>(36.9) | 1.73–9.36<br>(4.70) |
| *Skomer* | 5<br>(20) | 5,748–198,796<br>(49,789) | 38–260<br>(127) | 2.66–92.1<br>(39.9) | 13<br>(15) | 14,038–135,838<br>(75,343) | 30–281<br>(123) | 5.45–50.1<br>(12.1) |
| *Sule Skerry* | 2<br>(9) | 5,447–27,778<br>(16,612) | 101–152<br>(127) | 3.39–84.6<br>(44.0) | 7<br>(9) | 23,076–110,602<br>(71,731) | 44–221<br>(145) | 1.78–13.1<br>(7.53) |
| *Skokholm* | 0<br>(4) | N/A | N/A | N/A | 0<br>(0) | N/A | N/A | N/A |

**Supplementary Table 2.**
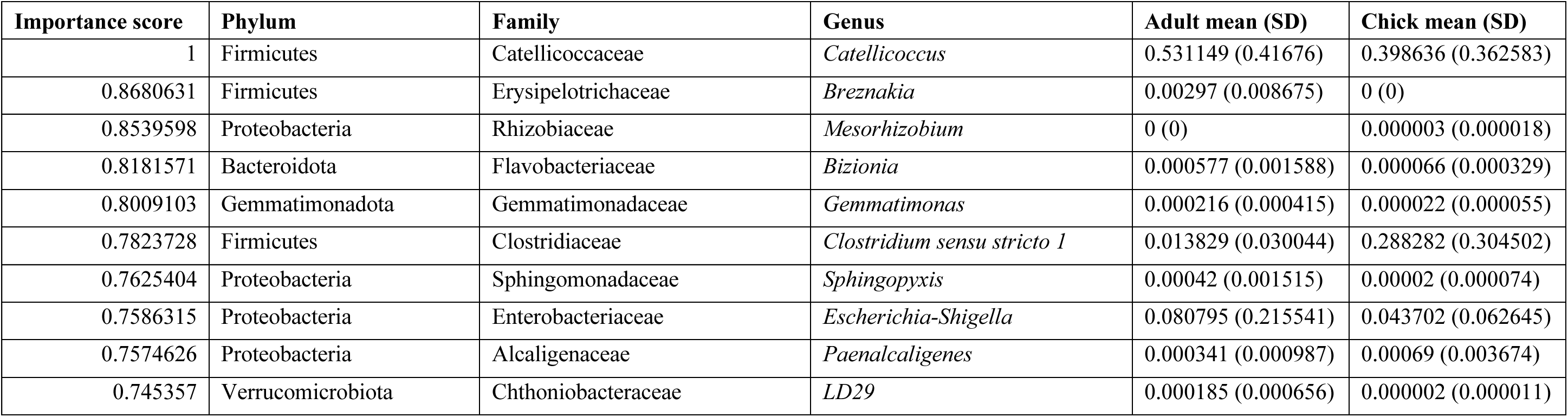
The top 10 genera with the highest importance scores in driving the age effect on puffin microbiota, based on a ‘drop one-taxon-out’ approach (see Methods). Importance scores reflect the extent to which excluding each genus increased uncertainty in the age effect estimate. Values for adults and chicks refer to mean relative abundance (on 0–1 scale) ± standard deviation across individuals within each age group.

